# Dream content and slow waves benefit preys against predators in a video game confrontation

**DOI:** 10.1101/2025.09.20.674534

**Authors:** Daniel S. Brandão, Rafael N. B. Scott, Ernesto S. Soares, Natália B. Mota, Sidarta Ribeiro

**Affiliations:** Brain Institute, Federal University of Rio Grande do Norte, Natal, Brazil; Bioinformatics Multidisciplinary Environment (BioME), Digital Metropole Institute, Federal University of Rio Grande do Norte, Natal, Brazil; Coimbra Institute for Biomedical Imaging and Translational Research, University of Coimbra (CIBIT), Coimbra, Portugal; Research Department, Mobile Brain, Rio de Janeiro, Brazil; Institute of Psychiatry (IPUB), Federal University of Rio de Janeiro (UFRJ), Rio de Janeiro, Brazil; Center for Strategic Studies, Oswaldo Cruz Foundation (FIOCRUZ), Rio de Janeiro, Brazil

**Author notes:** These authors contributed equally (Daniel S. Brandão, Rafael N. B. Scott). Corresponding authors (D.S.B); (S.R).

**Keywords:** Sleep, dreams, EEG, slow waves, prey, predator

## Abstract

Every studied animal species exhibits some form of sleep, a state evolved under the sustained influence of prey and predator behaviors. Dreams have been proposed to have evolved as offline mental simulations able to warn against impending threats from the environment, such as predators. To assess how the roles of prey and predator interact with sleep and dreaming, we used a first-person shooter game as a proxy for predator-prey confrontations. Electroencephalographic (EEG) and electrocardiographic (ECG) signals were recorded from 30 human adults paired and simultaneously recorded while 1) playing a video game round against each other, in which one participant was armed with a gun and the other was not, 2) taking a nap, 3) reporting dreams and/or thoughts, and 4) playing another round with the same gun assignment. We found that the participants in the prey role were highly affected by sleep and dreaming. Their score gains were positively correlated with slow wave activity during the nap, with how much the dreams were related to the game, and with heart rate variability during the first round. In contrast, no significant correlations were found for participants in the predator role. The results suggest that slow wave activity and game-related dream content during sleep improve the post-sleep performance of individuals under stressful, prey-like situations.

## Introduction

Sleep promotes memory consolidation, reorganizing them into stable representations (*1*) that are fundamental for survival and likely evolved under strong selection pressure (*2*). Pioneering studies proposed that slow-wave sleep (SWS, also called N3 sleep) primarily benefits declarative memories, while rapid eye movement (REM) sleep mostly helps non-declarative memory consolidation, such as procedural and emotional memories (*3*). However, later studies questioned this sharp functional division, showing for example that SWS promotes explicit knowledge of a motor sequence, highlighting its influence on non-declarative memories (*4*). The importance of the N2 sleep stage for the consolidation of a motor sequence has also been demonstrated (*5*).

Recent evidence shows that the immediate effects of the consolidation of skills and schemas are related to non-REM (NREM) sleep stages, and slower effects are related to REM sleep, which are better expressed after days or weeks (*6*). In addition, there is evidence of synergy between SWS and REM sleep, where optimal consolidation depends on the occurrence of these stages in succession (*7*), which corroborates the “sequential hypothesis” for sleep-dependent memory consolidation (*8*). Currently, it is conceived that SWS consolidates newly acquired memories by strengthening its neural representations, which are integrated with pre-existing memories during subsequent REM sleep (*9*).

There is also evidence that both NREM and REM sleep are fundamental for creative problem-solving (*10*). However, the influence of these stages depends on the nature of the proposed task. For example, anagram resolution (*11*) and the priming effect of words with weak semantic relation (*12*) are increased when participants are awakened from a REM sleep episode, in comparison with other sleep stages. These results point to the hyper associative character of REM sleep and suggest an explanation for the fact that dreams reported upon waking up during REM sleep are more bizarre than those of other stages (*13*).

The cognitive importance of SWS has been highlighted by some studies involving electronic games. In one study, the participants had to solve a puzzle until spending 10 min without being able to pass to the next level (*14*). At this point, part of the participants slept for 90 min and another part remained awake for the same time. After this period, the two groups were tested again at the same level they had previously failed to complete. Among the participants in the sleep group, 86% were able to resolve the level, while only 47% of those in the waking group were successful, a significant difference. A follow-up study found that participants who spent more time in SWS were more likely to solve the problem (*15*).

Brain activity patterns associated with different sleep stages have been proposed as mechanisms underlying memory consolidation and problem-solving. The benefits of SWS for memory have been associated with slow-wave activity (SWA) (*16*), and total time in SWS (*17*). In addition, some studies have shown that sleep spindles have a positive effect on memory consolidation (*18, 19*). Although these oscillations occur during both N2 and SWS, spindle characteristics such as density and amplitude differ between these stages (*19*). While some studies point to the importance of N2 spindles for the learning of motor sequences (*5*), other studies stress the role of SWS spindles in the consolidation of declarative memories (*20*). In addition, the temporal coupling between spindles and slow waves is strongly associated with memory consolidation (*21, 22*).

Another sleep-dependent mechanism linked to problem-solving is memory reactivation, where brain patterns associated with waking experiences are manifested again during sleep. This phenomenon has been observed most frequently in SWS, primarily in rodents (*23*) but also in humans (*24*). However, reactivation can also occur during REM sleep (*25*). The reactivation of memories during NREM sleep enables the abstraction of general rules from the information learned, while REM sleep favors the formation of new associations in a creative way (*10*).

Some theories speculate that the reactivation of memories may be the mechanism associated with the occurrence of dreams related to tasks performed during wakefulness (*26*). Because the chance of reporting a dream was significantly higher upon awakening from the REM stage than from the NREM stage, the idea that REM sleep was equivalent to dreaming initially became dominant (*27*). However, this idea began to be deconstructed with the improvement of the methodology for the study of dreams, resulting in the inclusion of a greater number of reports of mental activity obtained upon awakening during NREM, within the definition of dreams, identifying the possibility of dreams occurring during both stages of sleep (*28*).

One of the most interesting theories about the functions of dreams is the Threat Simulation Theory (TST), which argues that dreams are a biological mechanism evolutionarily selected as an offline state susceptible to safe training against real threats (*29*). The theory brings several predictions grouped into six propositions, which have already been corroborated by several studies. For example, it has been shown that dreams present threats more frequently and severely than real life, that dream threats are realistic, and that they mainly threaten the dreamer himself, who tends to react defensively (*30*). In addition, studies show that the dreams of traumatized children in war regions present more threats than the dreams of children without as much trauma or who live in quieter places (*31*). TST has been criticized, mainly by the fact that a small percentage of dreams involves realistic threats and when these occur, the dreamers rarely obtain success (*32*). However, the scarcity of threats in dreams can be a consequence of the safety promoted by modern society, as suggested by studies about the dreams of traumatized individuals (*31*). Corroborating this idea, during the survival pressure promoted by the Covid-19 pandemic, dreams became more related to “contamination” and “cleanness” (*33*). Altogether, this body of evidence shows that TST is a promising theoretical framework for understanding dream function (*34*), and requires further investigation.

One of the TST’s six propositions is that “perceptually and behaviorally realistic rehearsal of any skills, in this case threat-avoidance skills, leads to enhanced performance regardless of whether the training episodes are explicitly remembered”. The perception of the occurrence of this rehearsal during sleep independent of the participant’s recall would involve the detection of memory reactivations, a technique that still requires much improvement. However, regarding the remembered rehearsals, this proposition has been considerably investigated through research involving the analysis of dream reports. This type of survey involves performing a task before and after a sleep opportunity. In 2009, TST proponents stated that “The relationship between threat simulation and performance (speed and accuracy) could be investigated in the future by exposing participants to severe threats in a virtual reality environment or immersive video game and studying the dream rehearsal rates and performance in relation to each other” (*34*).

To evaluate these open questions, we conducted experiments involving the recording of electroencephalographic (EEG) and electrocardiographic (ECG) signals from pairs of human participants during rounds of a first-person shooter (FPS) video game confrontation in which one participant played against the other, simulating a realistic battle for survival (Fig. 1). In this game, participants assumed contrasting roles that simulated a predator-prey scenario. This dynamic is promoted by granting the predator a major competitive advantage (a gun), in comparison with the prey (no gun). After the first round, both participants slept in the laboratory, and then held a new round of dispute.

**Fig. 1.**
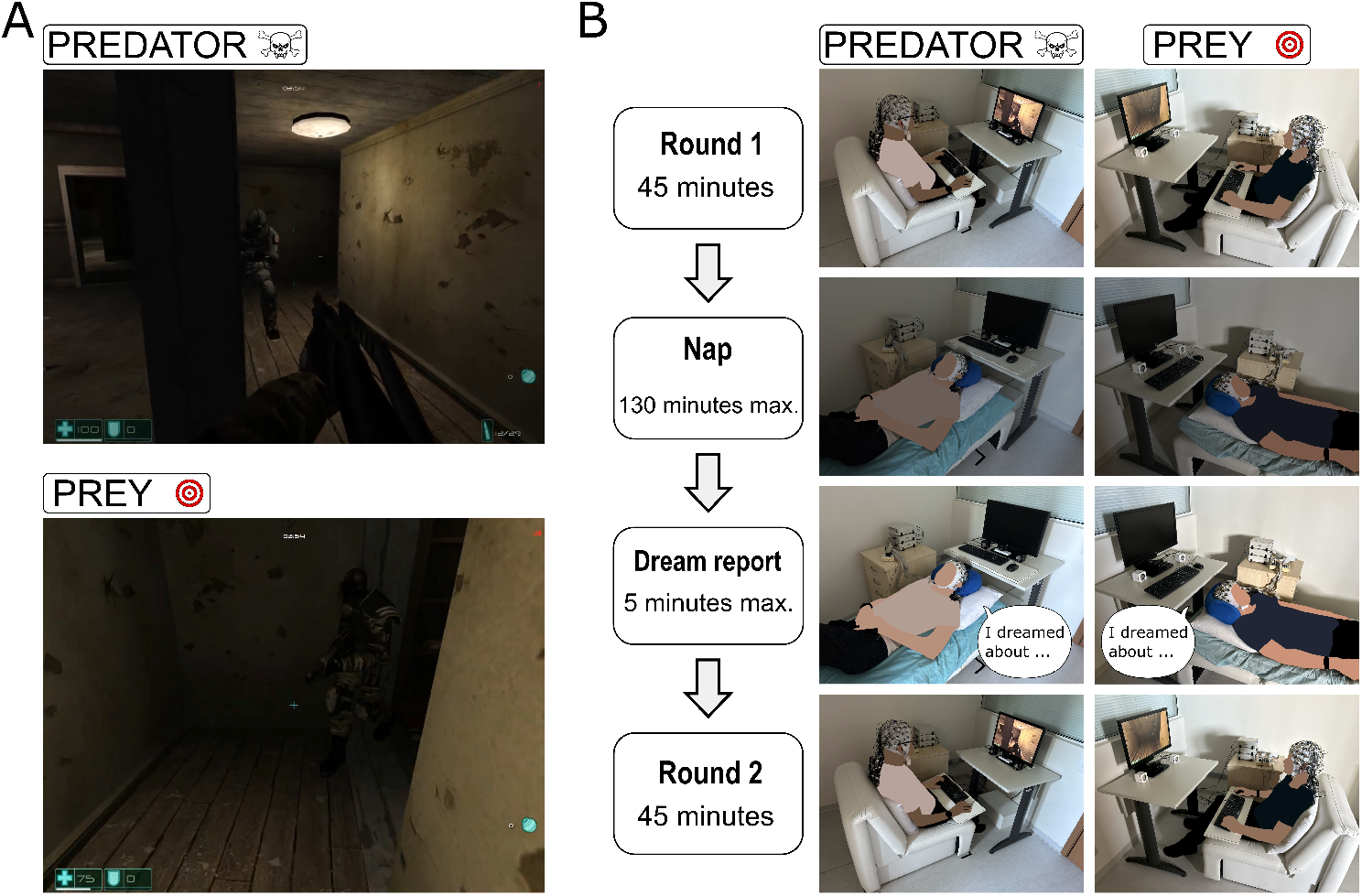
Experimental design. (**A**) Example of in-game screenshots for predator and prey participants. (**B**) Graphical description of the experimental procedure: Two participants go to the laboratory and are randomly allocated to Prey or Predator roles. Then, they play a first-person shooter game against each other for 45 min, where the Predator hunts the Prey. Next, both participants have a nap opportunity for up to 130 min, followed by oral description of any dream experiences or thoughts during the nap period. Finally, the participants played another round of the game, in the same prey-predator roles as before.

## Results

### Sample demographics

Table S1 presents the comparison of variables related to gender, experience in games, age and sleep patterns of the participants in the predator and prey group. No significant differences were found. Table S2 shows a comparison of sociodemographic, video game and sleep characteristics of the two data sets used in our study (morning and afternoon recordings). As no significant differences were found between the subsets, they were pooled into a single data sample as presented henceforth.

### Game scores

Score comparisons across the 2 rounds were performed for all participants with valid electrophysiological signals (N=13 for both predator and prey groups). 4 participants were included without the respective adversaries, when only one participant per pair had good-quality EEG/ECG recordings (see the Participants section). Significant score differences were not found between the prey and predator groups, neither for round 1 (adjusted p = 0.4222) nor for round 2 (adjusted p = 0.4236) (Fig. 2A,B). However, when separately evaluating the events of the game (wins, losses, and collections), a pattern was found in round 1 that was repeated in round 2, where the predator had more wins than the prey (adjusted p < 0.0050), in addition to having fewer losses (adjusted p < 0.0065) and fewer collections (adjusted p < 0.0011) (Fig. 2A,B). Thus, the experimental design successfully simulated a confrontation in which the predator had a survival advantage and the prey had greater success in collection.

**Fig. 2.**
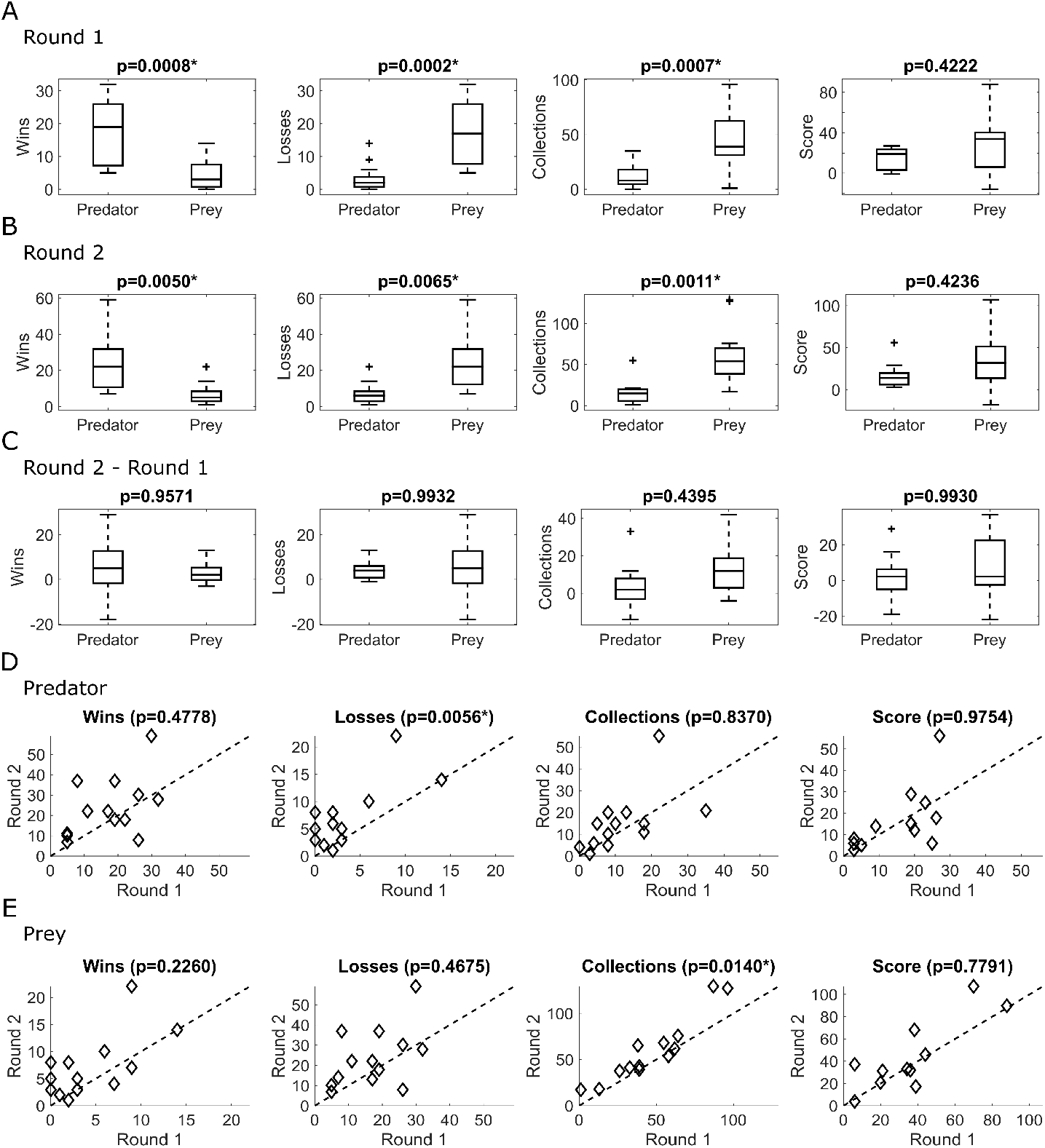
The video game simulation effectively influenced prey and predator behaviors, and the performance gain on round 2 is better seen for preys. The comparison between roles is being shown for **(A)** Round 1, **(B)** Round 2, and **(C)** Gain (Round 2 – Round 1). The evaluation to compare rounds is presented for **(D)** predators, and **(E)** preys.

Also, the score gains (round 2 minus round 1) were compared between predators and preys (Fig. 2C). No significant differences were found for the gain in the number of wins (adjusted p = 0.9571), losses (adjusted p = 0.9932), collections (adjusted p = 0.4395), and the overall score (p = 0.9930).

It was also evaluated whether prey and predators performed better in round 2 than in round 1 in relation to overall score, number of wins, losses, and collections (Fig. 2D,E). Regarding the overall score, no significant difference was found between the two rounds, both for the prey and the predator (adjusted p > 0.7791). A significant difference across rounds was found for the number of predator losses (adjusted p = 0.0056) and for the number of prey collections (adjusted p = 0.0140).

### Reports of mental activity

There was good consistency between the raters of the reports for the question that evaluates whether the participant dreamed, based on the Intraclass Correlation Coefficient (ICC=0.89). In relation to the questions assessing the characteristics of dreams, the ICC for the question that evaluates whether the participant clearly remembered the dream was moderate (ICC=0.66), while for the other questions it was good (ICC range: 0.76 to 0.84). The correlation analyses between dream properties and score gains revealed a significant positive correlation between game-related dreaming and score gains for participants in the prey role (Table 1 and Fig. 3, p = 0.013). None of the remaining dream properties showed significant correlations among prey participants. Furthermore, predator participants showed no significant correlations between dream properties and score gains (Table 1). Representative examples of dream reports with high and low relationship to the game are presented in the Supplementary Text.

**Table 1.** Correlations between dream properties and score gains.

| Role | Dream property | N | Rho | Adjusted p value |
| --- | --- | --- | --- | --- |
| Predator | Dreamed | 14 | -0.1781 | 0.5308 |
| Predator | Remembers clearly | 6 | -0.6667 | 0.5222 |
| Predator | About game | 6 | -0.6375 | 0.5806 |
| Predator | About laboratory | 6 | -0.2125 | 0.9861 |
| Predator | About personal life | 6 | 0.4928 | 0.7778 |
| Predator | About being prey | 6 | -0.2704 | 0.9778 |
| Predator | About being predator | 6 | -0.2704 | 0.9778 |
| Prey | Dreamed | 14 | -0.2619 | 0.3593 |
| Prey | Remembers clearly | 10 | -0.4970 | 0.5162 |
| **Prey** | **About game** | **10** | **0.8624** | **0.0136 \*** |
| Prey | About laboratory | 10 | 0.6335 | 0.2420 |
| Prey | About personal life | 10 | -0.7660 | 0.0702 |
| Prey | About being prey | 10 | 0.3620 | 0.8225 |
| Prey | About being predator | 10 | 0.3158 | 0.9042 |

**Fig. 3.**
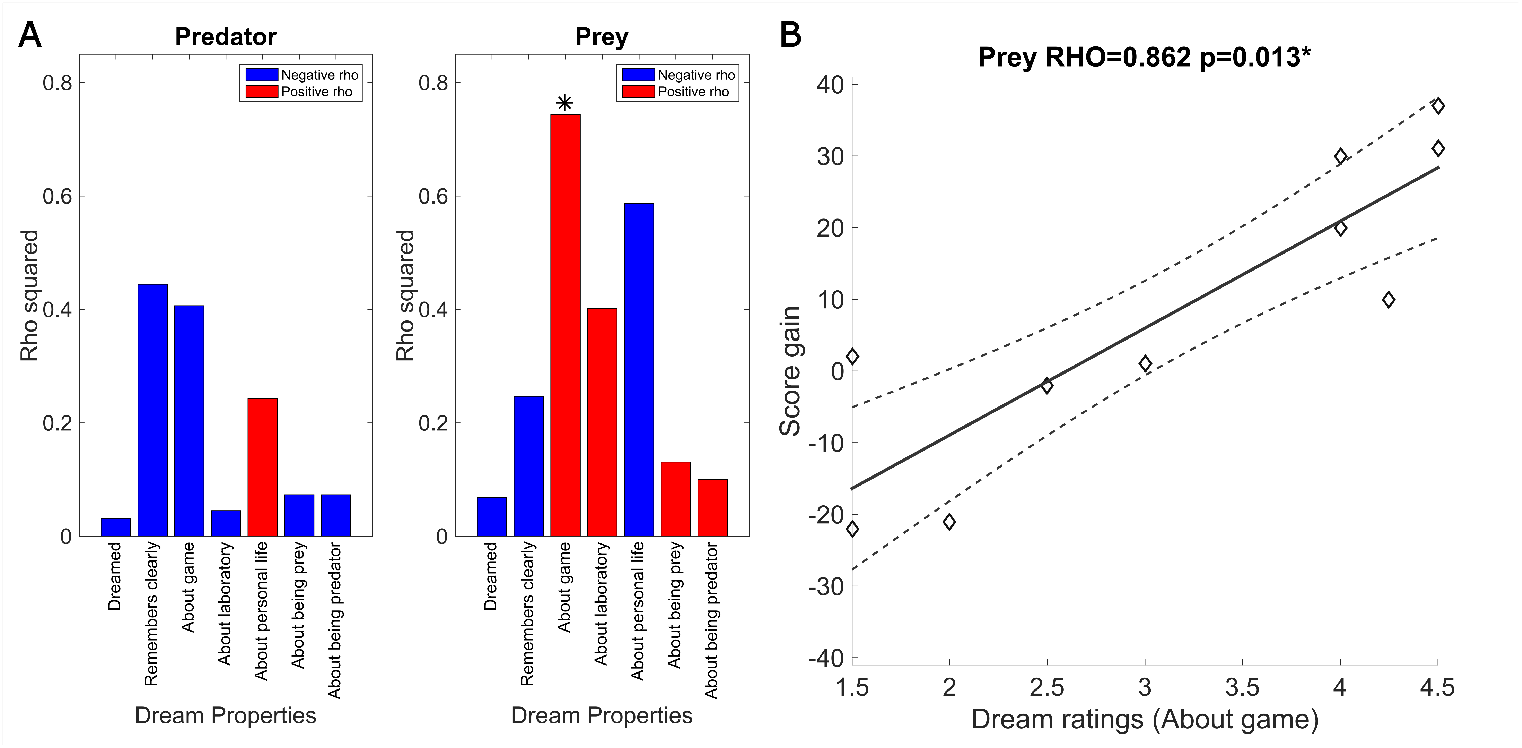
For participants in the prey role, dream reports about the game are associated with significant score gains. (**A**) Bar plots for Spearman correlation coefficients (Rho) of score gains and dream reports properties for predator and prey. (**B**) Scatter plot showing the correlation between score gain and ratings of the certainty that the dream is about the game.

### EEG Power across frequency bands

The correlation between EEG power in each frequency band of interest and the score gain in the game was verified through the topographic distribution of the correlation coefficients (Fig. 4A). Significant correlations occurred for the delta and beta bands for the prey group (Table 2). In delta, positive correlations were found in spatially diffuse channels, with a lower p-value in channel C2 (adjusted p = 0.0134) (Fig. 4B). In the beta band, a negative correlation was observed in the C4 channel (adjusted p = 0.0416) (Fig. 4C).

**Table 2.**
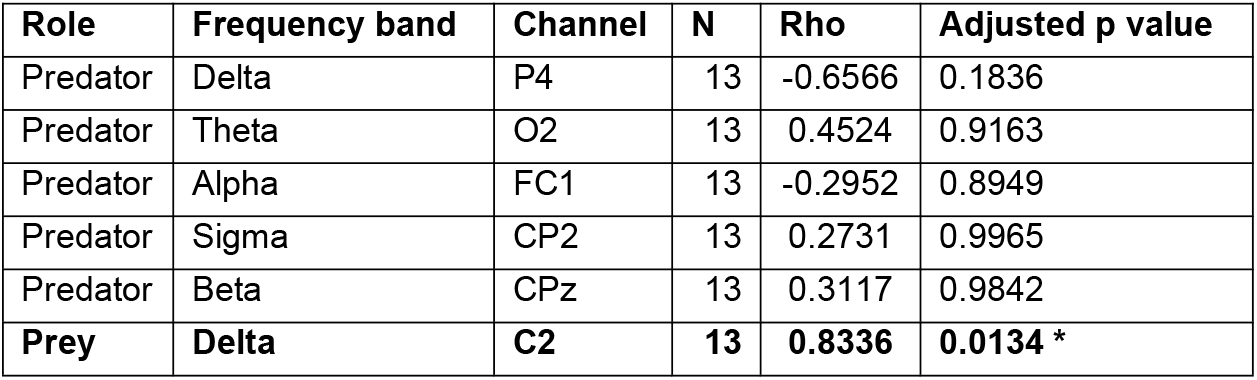
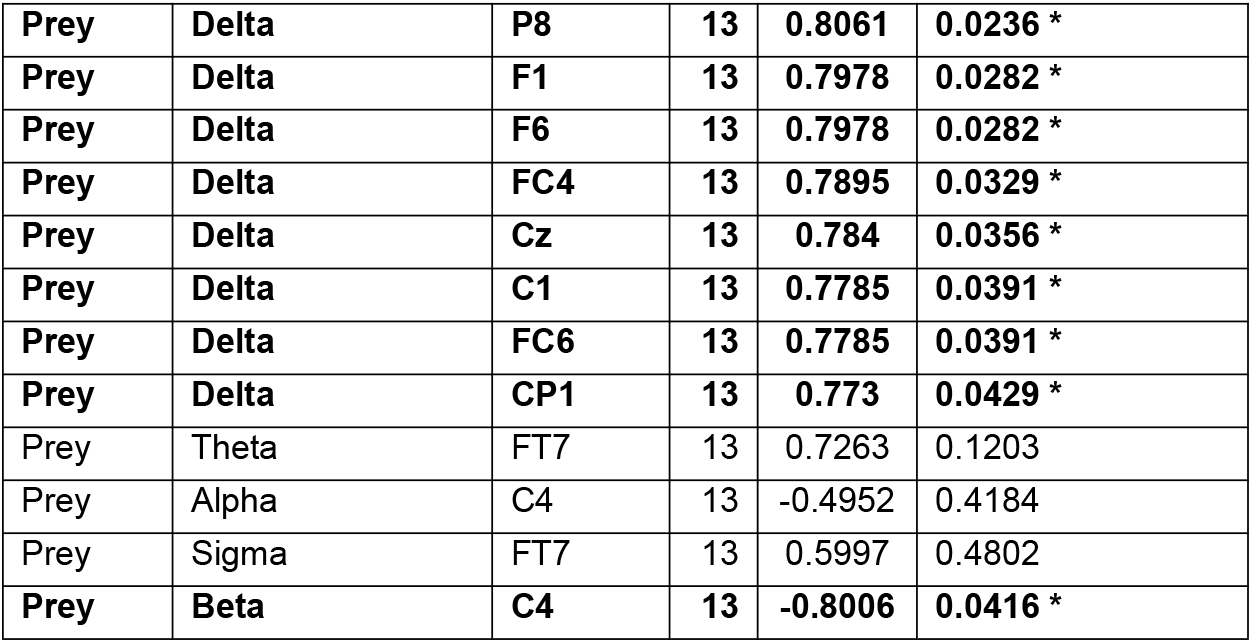
Correlations between score gains and EEG spectral power during the nap. For every combination of frequency band and role, the results for the channel with smallest p-value is being reported, including all channels with significant correlations.

**Fig. 4.**
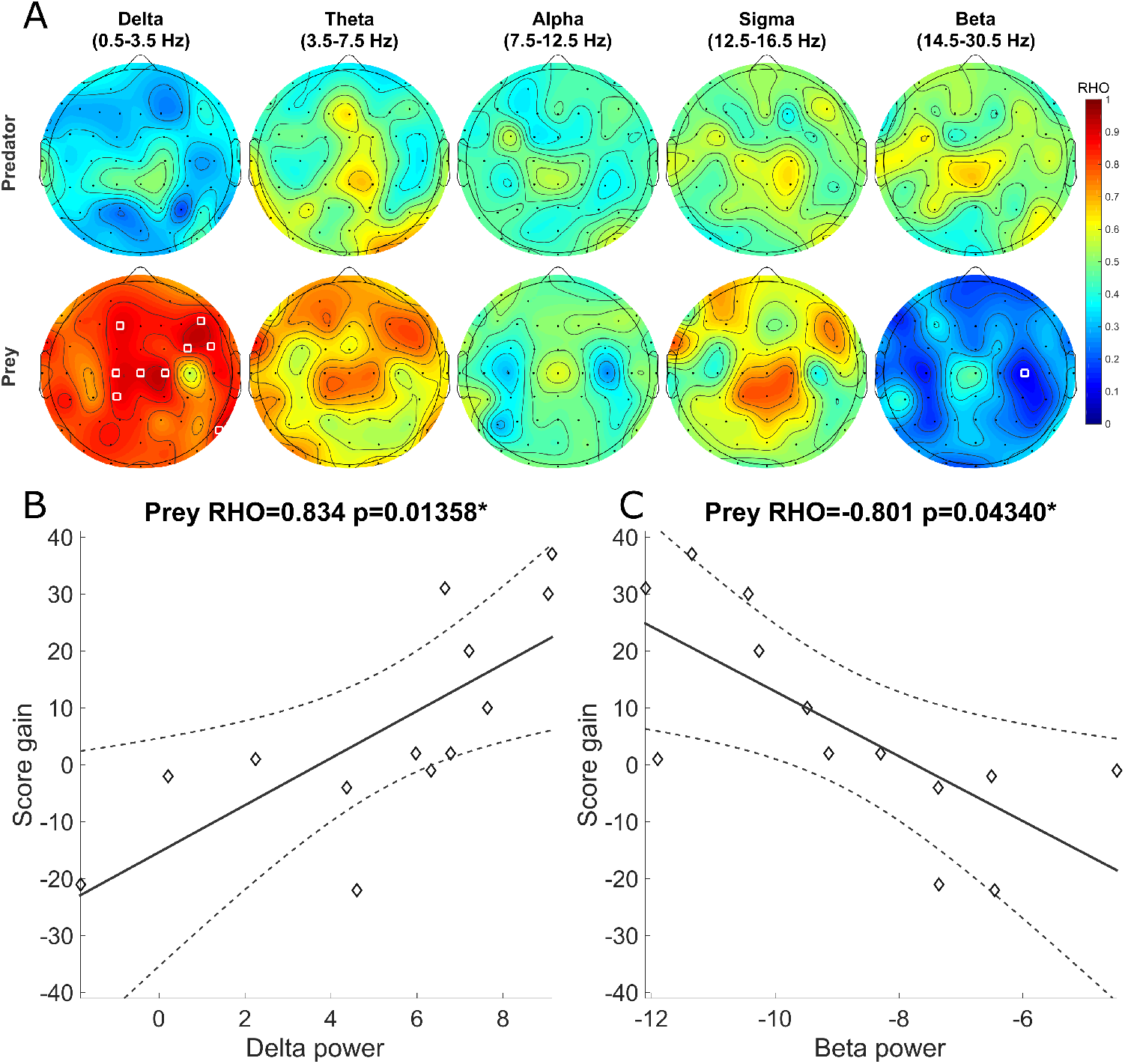
In the prey but not in the predator group, delta and beta power during sleep showed positive and negative correlations with score gains, respectively. (**A**) Topographies of Spearman correlation coefficients (Rho) between score gains and EEG spectral power during sleep. (**B** and **C**) Scatter plots showing the correlation between score gain and delta power at channel C2 (B) and beta power at channel C4 (C) for prey participants.

### Slow waves

Next, we evaluated whether score gains were correlated with slow wave properties during the intervening nap, specifically, the average amplitude, density, and summation of amplitude, in either preys or predators (Fig. 5A). Positive correlations were found for the summation of amplitude (Table 3), with lower p-value for channel FC1 (adjusted p = 0.0115) (Fig. 5B).

**Table 3.** Correlations between score gains and EEG slow wave parameters. For every combination of slow wave property and role, the results for the channel with smallest p-value is being reported, including all channels with significant correlations.

| Role | Slow wave properties | Channel | N | Rho | Adjusted p value |
| --- | --- | --- | --- | --- | --- |
| Predator | Mean amplitude | P4 | 13 | -0.6152 | 0.2648 |
| Predator | Density | PO7 | 13 | 0.309 | 0.9638 |
| Predator | Summation | POz | 13 | -0.5628 | 0.3223 |
| Prey | Mean amplitude | FC6 | 13 | 0.696 | 0.1128 |
| Prey | Density | CPz | 13 | 0.5557 | 0.4759 |
| **Prey** | **Summation** | **FC1** | **13** | **0.8336** | **0.0118 \*** |
| **Prey** | **Summation** | **F6** | **13** | **0.773** | **0.0363 \*** |
| **Prey** | **Summation** | **F1** | **13** | **0.7675** | **0.0390 \*** |

**Fig. 5.**
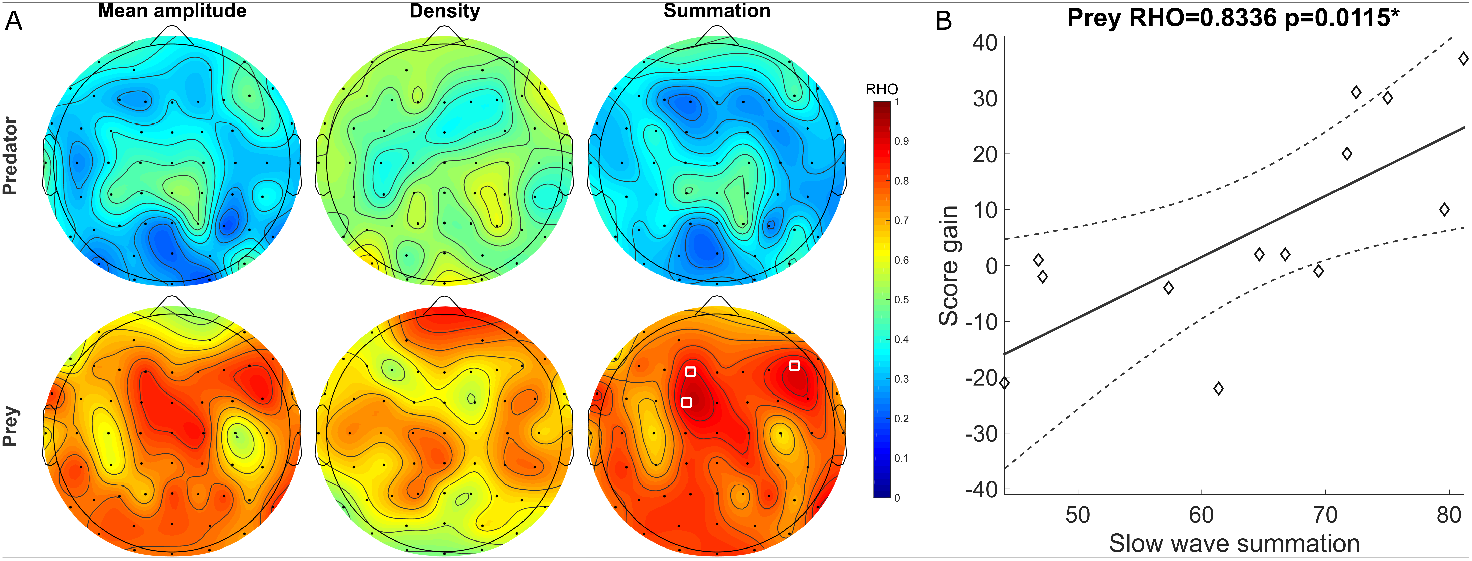
Slow wave summation was correlated with score gains in the prey group, but not in the predator group. (**A**) Topographies of Spearman correlation coefficients (Rho) between score gains and slow wave properties during sleep. White circles indicate significant electrodes. (**B**) Scatter plot showing the correlation between score gain and the summation of detected slow wave amplitudes at channel FC1 during sleep.

### Sleep spindles

At first, our intention was to compare the results obtained when evaluating the sleep spindles recorded during stage N2 with those of SWS. However, in SWS, sleep spindles were found in only 7 predators, showing no significant correlation with score gains. For the prey, only 2 participants had some sleep spindles in SWS, making it impossible to perform the analysis for this role. The number of sleep spindles found during stage N2 was higher and, for this reason, only the results for sleep spindles identified during N2 are being reported.

Next, we evaluated the correlation between score gains and the number or duration of sleep spindles (Fig. 6A), but no significant correlation was observed (Table 4). The correlation with lowest p-value was detected for the FC6 channel (adjusted p = 0.1549, non-significant; Fig. 6B).

**Table 4.** Statistics on spindle properties. Results for correlations between score gains and EEG spindle parameters

| Role | Spindle property | Channel | N | Rho | Adjusted p value |
| --- | --- | --- | --- | --- | --- |
| Predator | Spindle count | PO7 | 12 | -0.5579 | 0.4205 |
| Predator | Spindle duration | T7 | 12 | 0.5228 | 0.7936 |
| Prey | Spindle count | FC6 | 10 | 0.5636 | 0.4354 |
| Prey | Spindle duration | FC6 | 10 | 0.7455 | 0.2234 |

**Fig. 6.**
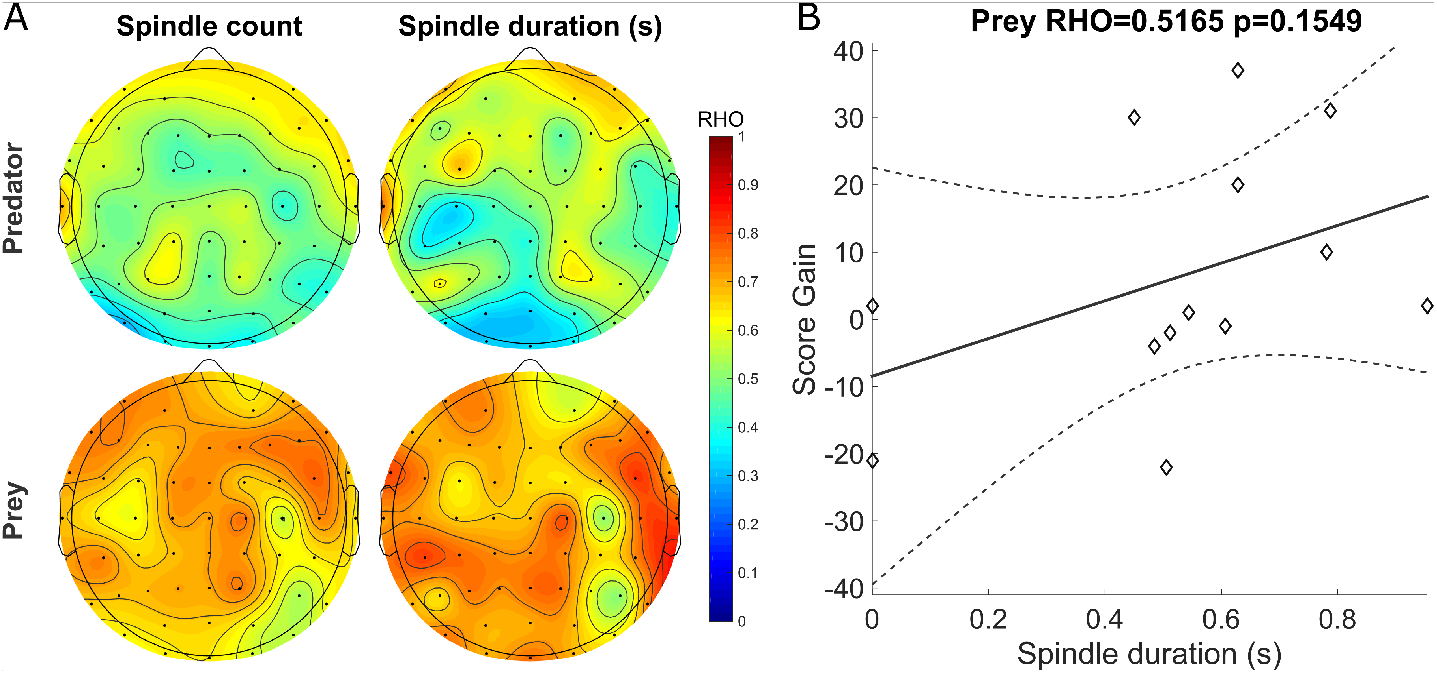
No significant correlations were found between score gains and spindle properties. (**A**) Topographies of Spearman correlation coefficients (Rho) between score gains and spindle properties. (**B**) Scatter plot showing the correlation between score gain and the spindle duration at channel FC6 during sleep.

### Sleep stages, sleep scales, and heart rate metrics

Out of the 26 participants, none woke up from SWS, 12 woke up from N2, 6 from N1, 5 from REM and 3 were already awake. The percentages of time elapsed in each stage occurred according to the following distribution (mean ± standard deviation [minimum – maximum]): SWS (3.38 ± 5.70 [0-18.75]), N2 (19.86 ± 31.30 [0-61.60]), N1 (7.03 ± 10.76 [0-28.64]), REM (11.12 ± 7.01 [0-53.12]), Awake (31.43 ± 47.53 [3.57-100]). The duration of the nap interval varied mainly due to the attempts to wake up participants from REM (111.86 ± 9.09 [94.04 – 130]).

For each stage, the number of people who did not spend any time in that stage was evaluated. Among the 13 preys, these quantities were 9 for SWS, 2 for N2, 3 for N1, 6 for REM and 0 for Wakefulness. The amounts for the 13 predators were 6 for SWS, 1 for N2, 1 for N1, 4 for REM and 0 for Wakefulness.

There were no significant correlations between score gains and the time spent in each sleep stage (adjusted p > 0.2645), the values of the sleep scales (adjusted p > 0.2473) or the heart rate metrics (adjusted p > 0.1399), while correcting for number of tests performed in each of these 3 analyses.

However, while evaluating only the variables selected for stress and sleep quality (see Materials and Methods), a significant correlation was found for the standard deviation of heart rate (sdHR) (N=8; rho=0.7857; adjusted p = 0.0406), but not for the PSQI (N=8; rho=0.1707; adjusted p = 0.8849). A comparison between roles was also performed for these selected variables, but no significant difference was found for sdHR (p = 0.9088) or for PSQI (p = 0.5850).

## Discussion

Significant correlations were obtained in the prey group between score gains and the amplitude of delta oscillations during the intervening nap and with game-related dreaming. However, in the predator group, no significant correlations were found. The results corroborate previous studies that show sleep and dream benefits to task performance (*26, 35*–*37*). They also indicate that there are major differences depending on whether the participants play the prey or the predator role.

In the prey group, performance gains were positively correlated with EEG power in the delta band and negatively correlated with EEG power in the beta band. The topographic analysis shows that the delta association was widely distributed, and that the channels with significant effects were dispersed in the scalp. For the beta band, the effect was accentuated specifically in the central right region.

EEG power in the delta band during sleep derives from slow wave oscillations. For the analysis of the individually identified slow waves, a positive correlation was found between the gain in the score of the prey and the sum of the amplitude of the waves in each segment, pointing in the same direction as the results found for EEG power in the delta band. These results are mutually reinforcing, as they are similar measures that are both influenced by the average amplitude and the density of slow waves. However, no correlation was found with performance when these other factors were considered in isolation. The improvement in prey performance depends on the increase in slow wave amplitude combined with the amount of slow waves during sleep, which supports a cumulative mechanism coupling SWS and learning.

Slow waves during sleep usually have a greater amplitude in the frontal region (*38, 39*). Benefits in memory consolidation tend to be mediated by amplitude increases of the frontal slow oscillations (*16*). However, in the present study, the score gain for the prey was not correlated more intensely in the frontal regions, but rather in a distributed way across the scalp.

A possible explanation for this atypical pattern may be based on the influence of stress on these individuals. Pre-nap stress has been shown to promote a statistically significant decrease in slow-wave amplitude during the first 30 min after bedtime when assessing the frontal, central, and parietal regions (*40*). In addition, one study showed that slow wave amplitude and density, and time in SWS are all decreased both in response to a previous stressful event and in preparation for later stress (*41*). In the present study, the game was a stressor for the prey both before and after the nap, and the two game sessions contribute to the decrease in slow wave amplitude.

Stress before bed has also been linked to increases in beta waves during sleep. A study involving the collection of EEG and cortisol levels from 154 participants showed a significant correlation between cortisol levels and the potency of beta in central regions during sleep (*42*). Another study showed that beta waves also increase in response to a stressor event and that this increase occurs in both N2 and SWS and persists throughout the night (*43*).

However, delta power decrease and beta power increase are also correlates of decreased sleep quality. Several studies have shown that individuals with primary insomnia have these effects across the entire night (*44*), but especially during the onset of sleep (*45*–*47*). Increased beta is a more consistent marker of lower sleep quality than delta decrease (*48*–*50*). The predominance of this effect on sleep onset was demonstrated in a study evaluating sleep throughout the night, in which beta power in individuals with insomnia was higher than in the control group only when observing the first NREM period (*51*). In addition, studies in individuals with insomnia show that the increase in beta power and especially the decrease in delta power during NREM sleep are predictors of their perceived stress levels (*52, 53*). In sum, studies of stress and lower sleep quality show that their effects are concentrated in the early periods of sleep and that the experimental design involving a nap of at least 30 min is sufficient for the observation of these effects.

The analysis of sdHR and PSQI score provided a way to differentiate the effects of stress on round 1 and sleep quality, respectively, of the preys. The significant correlation found between sdHR and score gain further support the notion that the decrease in the amplitude of slow waves and the increase in beta waves found in the prey group are both caused by stress, since these effects are typical of studies involving stressful situations. We can infer that the stress perceived during the prey’s sleep promotes lower score gains and, therefore, the preys that perceived less stress obtained greater performance gains. Also, since stress reduces time in SWS (*41*), the stress induced by the game can explain the fact that 9 of the 13 preys did not spend any time in SWS. This indicates that even out of SWS, SWA can benefit prey performance.

Stress, sleep, and slow waves have a complex interaction with the consolidation of memories. Both stress and sleep promote an increase in consolidation (*54*). However, the action of sleep on memory is mediated by the increase in the amplitude of slow waves (*16*). Stress, on the other hand, seems to do the same, but by the opposite mechanism, reducing slow waves (*55*). This reasoning is strengthened by demonstrations that stress impairs neutral memories but strengthens emotional memories (*56, 57*).

Sleep and stress may act together to benefit the prey group, specifically consolidating the most relevant memories to promote performance gains for participants in this role. In contrast, task performance in the predator group, received no benefits of stress. We found no difference in the sdHR between prey and predator roles, so we cannot affirm that the former was more stressful than the latter. However, we found that the sdHR was correlated with score gains only for preys. So, we deduce that stress was adaptive for preys, focusing memory processing on relevant game-related aspects.

No correlation was found between the occurrence of sleep spindles and performance gains. This result is unexpected, considering the strong relationship established in previous studies between sleep spindles and the consolidation of declarative and non-declarative memories, while evaluating N2 and SWS combined (*18, 19*). It has been shown that spindle density is greater in SWS than in N2 (*20*), but this evaluation was impaired due to the small amount of sleep spindles found during SWS in our datasets. A possible explanation is that the density of sleep spindles has been reported to be significantly reduced after a stressful situation (*58*), which in the current study is mainly associated with the prey condition. As discussed previously, the decrease of time in SWS promoted by stress also contributes with the difficulty to find sleep spindles at that stage in the current study.

A sufficient number of spindles was detected for the analysis within N2, but for this stage no significant correlation was found in any channel, neither for prey nor for predators. This agrees with the fact that the influence of spindles on memory consolidation is specific to SWS (*20*) and that memory consolidation is correlated with the power of slow waves, but not of spindles, also showing a positive correlation for time in SWS and a negative correlation for time in N2 (*59*).

Furthermore, there is evidence that spindles only contribute to memory consolidation when followed by a REM sleep episode (*60*). Moreover, evidence indicates that the contribution of spindles for memory consolidation occurs specifically during the transitions of SWS to REM (*9, 61, 62*). As 4 of the 13 predators and 6 of the 13 preys had no time in REM sleep, a considerable portion of the participants remained deprived of the spindle-related benefits of transitioning from SWS to REM sleep. The absence of REM sleep in several participants can be explained by the fact that the experimental design established a 130 min nap period. Despite being longer than the 90 min commonly expected for the first occurrence of REM sleep, factors such as presence in the laboratory and task-induced stress result in a generalized increase in sleep latency (*40*).

The analysis of dream reports indicates that the score gain of the preys in the second round is positively associated to the occurrence of game-related dreams in the prior nap period. Dreaming about the game right after playing it is a phenomenon extensively reported in the literature (*63*–*65*). The promotion of performance gains through the occurrence of dreaming was demonstrated in studies involving maze navigation (*26, 66*). However, the factors that promote or hinder this performance gain are still unclear, since in another similar study, supposedly due to the mere inclusion of a time counter associated with a financial reward, dreams did not benefit the task (*67*).

One way to reconcile these results is to consider that the association with a financial gain brings into dreaming certain concerns that are not task related. This aligns well with the presence of a marginally significant negative correlation in the prey group between score gains and the occurrence of dreams related to the participant’s personal life (p = 0.0702). This suggests that dreaming by itself is not enough to improve performance; rather, for that, the dream must be related to the task.

It is interesting that the performance of preys was related both to slow waves and dream content. This may be the mechanism explaining why the effect of dreams on memory consolidation is bigger for dream reports collected during NREM, compared to REM (*37, 68*). Dream content was also evaluated in participants who took the role of predator, but no significant correlations were found. One possible explanation is that preys mostly relied on spatial learning of the environment, while predators benefited more from combat-related procedural skills. Previous studies have shown dream-related improvements in the procedural performance of virtual tasks such as flying (*36*), playing tennis (*35*), or pressing a key pattern (*69*), but none including a prey versus predator confrontation.

Some limitations constrain how much can be extrapolated from the present study. First, the design did not include a control waking group, which would allow us to compare the gains associated with game-related dreams with those associated with game-related mentation during waking (*69*). Second, research challenges related to the interruption of research during the COVID-19 pandemic resulted in the division of these participants into two datasets (see Methods). These datasets were similar but presented differences regarding starting time of the experiment and partial sleep deprivation. Another caveat was the predominance of male participants, which makes it impossible to generalize the findings to females.

Despite these limitations the present results add support to the threat simulation theory (*29, 34*), as they suggest that sleep and dreams are important tools for adaptation when people play the prey role, but not the predator role. Predators were already at an advantage in the dispute and possibly did not need adaptation, and/or because the playful situation of the video game is not enough to mobilize the dreams of the predators. Given that the prey role, as designed in this study, was capable of engaging dreams and slow waves, future studies shall clarify which aspects of the prey experience are better expressed in the human brain during sleep and dreams.

## Materials and Methods

### Participants

30 participants were recruited to the laboratory in pairs, forming a total of 15 experimental recordings of a prey vs. predator confrontation. The study was approved by the Ethics Committee of the Federal University of Rio Grande do Norte, opinion number 2753906. The recruitment of the participants was carried out through the social network of the Brain Institute of UFRN. The inclusion criterion consisted of being a healthy adult between 18 and 40 years old. Among the 30 participants of the experiment, only 5 were female. The exclusion criteria consisted of self-declarations of having sleep disorders, neurological, psychiatric, or psychological diseases. By these criteria, no participant was excluded.

Among the 30 participants, 3 had their EEG data lost due to failures in storage and backup. 1 participant had low-quality EEG signal after visual inspection. For these 4 cases, the EEG data of the other participant in the pair remained in good condition. In addition, in the pair of the first experiment, the dream reports of the 2 participants were not collected.

Therefore, the EEG and dream analyses had different samples, considering the set of participants who presented good-quality EEG or dream data. Thus, EEG analyses were performed for a sample of 26 participants (13 prey and 13 predators) and for the analysis of dream reports, the sample had 28 participants (14 prey and 14 predators). The intersection between these two samples consists of 24 participants, 3 of whom were women, aged 24.6 ± 4 [21 – 35] years (mean ± standard deviation [minimum – maximum]).

### Experimental design

The experimental session involved the simultaneous recording of two human participants engaged in a competitive video game confrontation for 45 min (round 1), followed by a nap opportunity of 90-130 min, and another 45 min video game confrontation (round 2) (Fig. 5). During all phases, EEG and other peripheral signals were recorded (see section Electrophysiological Data Acquisition).

To wake at least one of the participants in REM, the experimenters woke the subjects about 2 min after observing the first 30 s epoch of REM sleep in any participant. The duration range of the naps was chosen to increase the probability of REM sleep occurrence, since a complete sleep cycle including the REM stage lasts approximately 90 to 110 min (*70*).

First, a small study with 11 pairs of participants (dataset 1) was planned, which would be followed by a second study with more participants. In the second study, some methodological changes were made in relation to the first, seeking to improve the experiment. We conducted experiments from the second study with 4 pairs (dataset 2) and the research was interrupted by the COVID-19 pandemic. When we returned to activities, we could no longer perform two EEG recordings simultaneously because some electrodes showed intense dryness in the insulating covering, and we had financial problems for buying new electrodes. Thus, we decided to combine the two samples, considering that the main methodological aspects are preserved between the two samples. The overall methodology was preserved between the two datasets, despite a few differences. The experiments of dataset 1 were carried out in the afternoon (starting at 1:00 p.m.), while those of dataset 2 were carried out in the morning (starting at 7:00 a.m.). Each experimental session was about 6 h long. For afternoon recordings, the strategy was to facilitate naps due to post-prandial sleepiness. For the morning recordings, the participants were instructed to, in the previous night, sleep 2 h less than they were used to, thus promoting the sleepiness necessary for the experiment.

While the electrodes were being placed, the participants answered the following questionnaires: Pittsburgh Sleep Quality Index – PSQI (*71*), Fletcher and Luckett’s Questionnaire (*72*), Epworth sleepiness scale (*73*), in addition to questionnaires to assess aspects of the participants such as the sociodemographic context, sleep and activities of the day before the experiment and previous experience with electronic games.

The audio and video of the computer that was running the game were recorded through the software Fraps (https://fraps.com/) for dataset 1 and OBS Studio (https://obsproject.com/) for dataset 2. After each game round, a brief questionnaire was applied where each participant evaluated their performance during the game.

The participants were instructed to take a nap between the two rounds. After the nap, a report of mental activity was requested, where the participants reported on their dreams or thoughts immediately before being woken up. For dataset 1 a written report was requested, while for dataset 2 an oral report was recorded. For dataset 2, participants observed the elapsed recording time and were instructed to present a report of no more than 5 min. After producing the mentation report the participants answered a questionnaire to assess the quality of sleep in the laboratory.

### Video game

During rounds 1 and 2 the participants played against each other using the game F.E.A.R. Combat, a version of the game F.E.A.R. containing only multiplayer mode and available for free on the Internet (https://fearcombat.org/). The scenario used was “Residential Evil”, made available in an online community of game add-on creators (https://www.moddb.com/games/fear/addons/residential-evil). For the experiment, the game was set up to make one of the participants play in the role of a prey, while the other participant played in the role of a predator. Participants in the predator role were told that their goal was to kill the prey and to not be killed, while participants in the prey role were told that their goal was to collect medkits scattered around the environment while avoiding being killed. The participants had no information about the opponent’s goal. Both could attack the opponent with short-range melee attacks, but the predator had the advantage of also being able to fire a mid-range shotgun. One punch from either participant was enough to kill the opponent, while the shotgun could kill with one or more shots, depending on the shooting distance. The number of shots needed to kill the prey varies from 9 to 17 at long-range, 2 or 3 at mid-range and 1 at short-range. Despite needing several shots, the possibility of a mid-range kill gave the predator a major advantage over the prey. Thus, it was expected that the prey would tend to be killed in most encounters, and that this induced in them a fight-or-flight response.

### Game scores

For each round, based on the observation of the videos, the following variables were extracted: number of times the opponent was killed (wins); number of times the player died (losses); number of medkits collected (collections). The prey’s score = collections - losses, while the predator’s score = wins - losses. The main metric evaluated was the score gain = score_round2_ - score_round1_.

### Electrophysiological data acquisition

During all phases of the experiment (round 1, Nap and round 2) electrophysiological signals from EEG, EOG (electro-oculography), EMG (electromyography) and ECG were recorded through 64 channels. ActiCap electrodes, BrainAmp DC amplifiers and BrainVision Recorder recording software, version 1.20.0506, all manufactured by Brain Products, GmbH, were used.

The EEG electrodes were positioned in caps suitable for the size of the participants’ heads according to the 10–10 system (*74*). During recording, the electrical reference was positioned at FCz and the ground electrode at AFz. The sampling rate was set to 1000 Hz. The impedance of the electrodes was kept below 20 kΩ.

The 64 channels were positioned differently between the two samples. This occurred because in sample 2, another set of caps and electrodes was used, with different electrode positioning. However, 54 channels were in common between the two samples. During signal processing, the reference was changed to the average of all electrodes, turning the signal from the FCz electrode into a valid channel. Thus, the EEG analyses of this study are based on the following 55 channels: AF7, AF3, AF4, AF8, F7, F5, F3, F1, Fz, F2, F4, F6, F8, FT7, FC5, FC3, FC1, FCz, FC2, FC4, FC6, FT8, T7, C5, C3, C1, Cz, C2, C4, C6, T8, CP5, CP3, CP1, CPz, CP2, CP4, CP6, P7, P5, P3, P1, Pz, P2, P4, P6, P8, PO7, PO3, POz, PO4, PO8, O1, Oz and O2.

### EEG signal processing

The EEG signals were all processed through automatic algorithms implemented in MATLAB (https://www.mathworks.com/products/matlab.html), using the EEGLAB toolbox (*75*). The processing sequence applied was similar to the HAPPE pipeline (*76*). After the application of HAPPE and the visual inspection of the signals obtained, it was verified the need to modify some of its steps and to update some methods to more recent approaches. The main differences involve modifications in the methods of automatic rejection of channels and intervals with artifacts, the replacement of the way of performing the automatic correction of ocular artifacts and the filtering of data at the end of processing. Although EEG recordings are also available during both rounds of gameplay, in this work the electroencephalographic analysis was focused on the signals during the nap.

The first step of processing was the removal of channels, so that only the 54 channels common to the 2 samples are left. The second step was to change the reference of the EEG signals to the average of these 54 channels, adding the original reference (FCz) to the set of channels, forming a total of 55 channels. The third step was the application of a bandpass filter between 1 and 250 Hz. Signals outside this frequency range impair the automatic removal of artifacts (*76*). The fourth step was the removal of electrical interference that occurs at 60 Hz, through a multi-taper regression using the EEGLAB plugin called CleanLine. This method makes it possible to remove electrical noise without distorting the EEG signal at neighboring frequencies, as traditionally happens with the use of notch filters.

The fifth step was the rejection of bad channels. This was done by evaluating the spectrum of the entire signal between 1 and 125 Hz, obtaining the logarithmic power in this band, averaging and standard deviation across all channels, and removing those that are 3 standard deviations above or below the mean (*76*). By visual inspection, it was verified that the evaluation of all bad channels needed to be performed twice per experimental block, because some bad channels are usually not identified the first time but are removed when applied a second time. Considering the two applications of the channel removal methods, together 4.88 ± 2.76 [0 – 12] channels were removed.

The sixth step was the removal of ocular artifacts through ICA (Independent Component Analysis). The HAPPE pipeline suggested the application of a wavelet-enhanced ICA (w-ICA), but we detected that this approach led to a big decrease in signal amplitude, compromising the spectral features of the signal. So, instead, we applied a traditional ICA. This method transforms the signals from a set of channels into a set of components. The signal from some types of interference, such as those related to eye movements, is usually restricted to a few of these components. By setting the weights of these components to zero and converting the signals back to the channel representation, these interferences are considerably attenuated. To identify the ocular components, we used ICLabel (*77*), which is an EEGLAB plugin that classifies ICA components into different types of artifacts. Specifically, ICLabel assesses the likelihood that each component originates from the brain, muscles, eyes, heart, electrical network noise, noisy channels, or another unknown source. Components that had a probability of ocular origin greater than 50% were removed.

The seventh step involved the removal of time intervals containing artifacts by applying two criteria based on amplitude and normalized power. For the amplitude criterion, HAPPE indicates the limit of 40 μV, but notes that this value is lower than usual due to the amplitude reduction promoted by w-ICA. Because we use traditional ICA, we adopted the 100 μV threshold, which is the value suggested for adult participants by the MADE pipeline (*78*). Thus, the points at which the absolute value of the amplitude (non-negative) was greater than this limit were marked as artifacts. In addition, for each point identified by this criterion, the points located 200 ms before and after were also marked as artifacts. The normalized power criterion was the same method used to remove channels but dividing the signal into 2-s epochs without overlapping, discarding epochs with power greater than 3 standard deviations from the average of the epochs of the channel.

The eighth stage involved a new removal of channels, specifically those that had more than 20% of the time identified as an artifact by the amplitude criterion. The criteria were then applied again to obtain the final identification on which intervals are artifacts. Evaluating the percentages of time identified, it was observed that by the amplitude criterion, an average of 12.73% were removed (standard deviation: 12.47%; maximum: 44.30%). On the other hand, through the normalized power criterion, an average of 12.55% (standard deviation: 3.01%; maximum: 20.20%) were removed. By combining the 2 criteria, an average of 18.47% were removed (standard deviation: 10.77%; maximum: 44.90%).

The ninth step involved interpolating the channels that were rejected in the fourth and eighth steps. For this purpose, the spherical spline interpolation method was used. This process consists of reconstructing the missing channels from the signal of all the other channels, where the closest channels have the greatest influence on the generation of the resulting signal.

The tenth step was the application of a 40 Hz low-pass filter. This step is not present in the HAPPE, which maintains the upper spectral limit at 250 Hz. However, frequencies above 40 Hz are already heavily contaminated by muscle artifacts (*79*). Thus, the application of this low-pass filter together with the removal of frequencies below 1 Hz performed in the second stage of processing, defines the spectral range between 1 and 40 Hz forming the set of frequencies present in the data resulting from this processing.

### Band power analysis

The relationship between the game scores and the power in different frequency bands during the block reserved for Nap was evaluated. This block was divided into 2-s segments and those that had artifacts in any channel or instant of time within the segment were discarded. The power spectral density was calculated for the valid segments, with a frequency resolution of 1 Hz. The values were then transformed to a logarithmic scale and represented in decibels.

The power bands evaluated were: Delta (1 to 3 Hz), Theta (4 to 7 Hz), Alpha (8 to 12 Hz), Sigma (13 to 16 Hz) and Beta (15 to 30 Hz). The sigma band is related to sleep spindles and the frequency range used was based on a study that showed the importance of the occurrence of spindles in this band for the consolidation of memories during sleep (*18*). The other bands, despite having their limits varying greatly between different studies, were defined according to a book chapter on computational EEG analysis (*80*).

### Wave pattern detection

In addition to spectral analysis, which evaluates the amplitude of oscillations generically within a time interval, algorithms were applied to identify individual occurrences of some wave patterns related to memory consolidation during sleep.

To identify each occurrence of sleep spindles, the A7 algorithm was applied, which can detect spindles with performance similar to that of experts in the field (*81*). For each channel, the number and average duration of the spindles occurred separately for the spindles that occurred during the N2 and SWS stages were evaluated. Participants who did not have any spindles identified in any channel were removed from the specific analysis of each stage. For participants in which the absence of spindles occurred only in some channels, the value 0 was assumed for the value of their properties.

For the identification of slow waves (delta), an algorithm similar to that described in a study on the effect of partial sleep deprivation on slow waves was implemented (*82*). The implementation differences involve changing the passband range from 0.5-4 Hz to 1-3 Hz, the stopband from 0.1-10 Hz to 0.1-3.9 Hz, and converting the signal to decibels (dB) to use the same parameters defined for delta-band analysis. The evaluation of the characteristics of the slow waves was carried out in 6-s epochs free of artifacts. For each channel, the average amplitude, the amount of slow waves and the summation of the amplitude (quantity * average amplitude) were obtained for each epoch. Then, the average of all epochs was made for each channel.

### ECG signal processing

The processing steps described below were performed independently for the ECG signal of each game round and for the nap. However, in the present study, only the results referring to round 1 were used, to provide variables capable of assessing the level of stress to which the participants were subjected before the nap.

The heartbeats of the ECG signal were identified using the Pan-Tompkins algorithm (*83*). Some subjects accidentally displaced or disconnected the ECG electrode during certain parts of the experiment and therefore these parts could not have their electrocardiographic measurements analyzed. Another drawback is that the movements of the participants during the experiment eventually caused artifacts in the ECG signal, which generate failures in the detection of the heartbeat through the automatic algorithm. Therefore, a script was developed in Matlab (Mathworks, Natick, MA) for the interactive inspection of these issues.

First, a visualization was generated containing 3 graphs, where one of them presented the heart rate in beats per min automatically obtained for each participant, highlighting the maximum and minimum values. The other 2 graphs showed the 2 intervals of the ECG that were associated with the minimum and maximum heart rate values. These graphs allowed the identification of artifacts, which result in the identification of false beats between real beats. By analyzing the signals in the intervals, using the interactive script, it was evaluated whether it was necessary to remove the interval between beats that associated the minimum or maximum.

When the removal was performed, the anterior and posterior intervals were also removed. At that moment, the visualization was updated to show the new minimum and maximum heart rate. This process was repeated until the extreme heart rate values were associated with ECG intervals that did not present errors in the identification of the heartbeat. The removal of intervals caused the introduction of continuity breaks in the heart rate, dividing the signal into several segments. To form a single, continuous signal, segments containing only 1 interval between beats were first removed. Then it was added 1 point between neighboring segments, which was calculated from a spline interpolation between the two segments.

After removing all extreme values containing artifacts, we used the HRVAS software (*84*), implemented in Matlab, to obtain variables related to statistical measures and heart rate variability. The correlation between these variables and the score gain of prey and predators was verified.

### Heart rate metrics

To measure stress during round 1, 11 heart rate metrics were calculated based on the interbeat interval (IBI), the heart rate (HR), and spectral aspects, like low-frequency (LF) and high-frequency (HF). The evaluated metrics were: range IBI (difference between maximum and minimum IBI values), mean IBI, median IBI, sdIBI (standard deviation of IBI values), RMSSD (root mean square of successive differences), mean HR, sdHR (standard deviation of heart rate), HRVTi (heart rate variability triangular index), nLF (normalized low-frequency power), nHF (normalized high-frequency power) and LFHF (the ratio of LF to HF) (*84*).

From these metrics, special interest is given to sdHR, as the main measure of stress during round 1, since it has been previously highlighted as the most significant heart rate metric to differentiate stress-inducing videos from non-inducing ones in a study evaluating participants while watching videos (*85*).

### Sleep scoring

Each 30-s epoch from the beginning of the record was automatically staged in 5 stages: Waking, REM, N1, N2 and SWS (N3). A method called CRNNeeg was used, which has a performance similar to that of experts (*86*). For scoring, this algorithm is based only on 4 channels, formed from the derivations of 8 electrodes: F3-C3, C3-O1, F4-C4, and C4-O2. From the scoring, the percentage of time in each stage was calculated, considering as total time the time between turning off and turning on the lights, defining the interval reserved for the nap opportunity.

### Sleep scales

For the sleep scales, the Epworth sleepiness scale, the total PSQI score, as well as its 7 subcomponents and the total sleep time of the last night were analyzed. As the main measure to assess sleep quality, the total PSQI score was selected, as it is a variable that consistently differentiates individuals with and without sleep disorders like insomnia (*48, 49, 52*).

### Evaluation of mental activity reports

The reports of mental activity obtained after the period reserved for the nap (possible dream reports) were typed from the written reports, in the case of Sample 1, and transcribed from the audio recordings, in the case of Sample 2. The reports were compiled in a spreadsheet and sent to 4 raters, who gave their opinions on each report for each of the following statements: 1 – The participant dreamed; 2 – The participant clearly remembers the dream; 3 – The dream was related to the game. 4 – The dream was related to the laboratory. 5 – The dream was related to the participant’s personal life. 6 – The dream represented the situation of being a prey. 7 – The dream represented the situation of being a predator.

The raters only read the reports and did not have access to other information about the participants, such as whether they were prey or predators, what their scores were in the game, age, gender, etc. The answers to the statements were recorded in the spreadsheet itself, which was sent back after complete filling. For each statement, the evaluators typed an “x” to identify their answers among the following options: Strongly disagree; Disagree; I disagree slightly; I agree slightly; Agree; I strongly agree.

In the columns of the spreadsheet, the options were presented in this same order, emphasizing the definition of a scale of agreement. For the evaluation of the answers, the number 1 was assigned to “Strongly disagree” and the number 6 to “Strongly agree”, with numbers 2 to 5 for the intermediate options. An option for neutral agreement was not included to force the choice between agreeing or disagreeing for each report, making it possible to convert the answers to a binary standard. The raters were instructed that, if they considered that the answer to statement 1, which evaluates whether the participant dreamed, was “I strongly disagree”, “I disagree” or “I slightly disagree”, the other statements about that report should be left blank.

For each report, the consensus of the evaluators was obtained from the calculation of the median of the raters for each statement. If the median of the answers to statement 1 of a report was greater than or equal to 3.5 (the middle of the agreement scale), it was considered a dream report. For the other statements that were left blank because the evaluator considered it not to be a dream report, the value of 3.5 was assumed as the answer.

To quantify the agreement between raters, the ICC was used. The values of this metric are being interpreted by the following guidelines: below 0.50: poor; between 0.50 and 0.75: moderate; between 0.75 and 0.90: good; above 0.90: excellent (*87*).

### Statistical analyses

In this study, Spearman’s correlation tests were applied to verify whether the game scores of both prey and predator were correlated with continuous variables that represent dream characteristics and wave pattern properties in EEG signals.

To control the increase in the rate of false positives promoted by the comparison of multiple variables, permutation tests were performed (*88*). These procedures perform the statistical test several times, shuffling the values of the variables being compared and verifying whether the statistical result obtained is greater than the chance of finding a significant result randomly for the values of the variables compared. For consistent p-value results up to 4 decimal places, it was found that 100,000 permutations were needed for the correlation tests. Thus, the p-values presented have already been adjusted to avoid problems of multiple comparisons.

To verify a priori differences between prey and predators, and between samples, the chi-square test was applied for categorical variables (sex, experience in games, etc.), and the Mann-Whitney U test for continuous variables (Pittsburgh and Epworth scales, etc.). For the a priori comparisons, statistical correction was not applied through the permutation test, to increase the chance of perceiving pre-existing differences, which may impair the interpretation of the data.

For the a priori comparisons, marginally significant results were observed, defined as those with p-value less than 0.1. In the other figures and tables, significant p-values are indicated as those with an adjusted p-value of less than 0.05, being represented by asterisks (*), except in topographies, where significant channels are identified by white squares.

For correlation plots, the solid line represents the simple linear regression and the dashed lines delimit the 95% confidence interval, indicating the region in which one can be 95% sure that it contains the mean of the values.

## Acknowledgments

We thank V.C. do Nascimento and L.M. Gila for helping in EEG data acquisition. We also want to thank the contribution of L.M. Gila, G.B.L.L. de Oliveira, J.L.L. Souza and E.A. Neves for rating the mental activity reports. We thank R.C. Moioli, D.B. de Araújo, D.Y. Takahashi, M.A.L. Miguel, F. Beijamini, J.G.O. Brockington, A.K.D.B. Filho, J.A. Queiroz, J.B.C. de Oliveira, T.F.P.M. Rodrigues and J.F. Araújo for insightful discussions.

## Author contributions

Conceptualization: D.S.B, R.N.B.S., and S.R. Methodology: D.S.B, R.N.B.S., and S.R. Software: D.S.B Formal analysis: D.S.B and R.N.B.S. Investigation: D.S.B, R.N.B.S, N.B.M., E.S.S. and S.R. Data curation: D.S.B. Writing—original draft: D.S.B. Writing—review and editing: D.S.B, R.N.B.S, N.B.M, E.S.S. and S.R. Visualization: D.S.B. Supervision: S.R. Funding acquisition: S.R.

## Data and material availability

The data that support the findings of this study are available from the corresponding authors upon reasonable request.

## References

1. S. Diekelmann, J. Born, The memory function of sleep. Nat. Rev. Neurosci. 11, 114–126 (2010).

2. A. C. Keene, E. R. Duboue, The origins and evolution of sleep. J. Exp. Biol. 221 (2018).

3. W. Plihal, J. Born, Effects of early and late nocturnal sleep on declarative and procedural memory. J Cogn Neurosci 9, 534–547 (1997).

4. J. N. Cousins, W. El-Deredy, L. M. Parkes, N. Hennies, P. A. Lewis, Cued Memory Reactivation during Slow-Wave Sleep Promotes Explicit Knowledge of a Motor Sequence. J. Neurosci. 34, 15870–15876 (2014).

5. S. Laventure, S. Fogel, O. Lungu, G. Albouy, P. Sévigny-Dupont, C. Vien, C. Sayour, J. Carrier, H. Benali, J. Doyon, NREM2 and Sleep Spindles Are Instrumental to the Consolidation of Motor Sequence Memories. PLOS Biol. 14, e1002429 (2016).

6. S. I. R. Pereira, P. A. Lewis, The differing roles of NREM and REM sleep in the slow enhancement of skills and schemas. Curr. Opin. Physiol. 15, 82–88 (2020).

7. S. A. Cairney, S. J. Durrant, R. Power, P. A. Lewis, Complementary roles of slow-wave sleep and rapid eye movement sleep in emotional memory consolidation. Cereb. Cortex 25, 1565–1575 (2015).

8. A. Giuditta, M. V. Ambrosini, P. Montagnese, P. Mandile, M. Cotugno, G. G. Zucconi, S. Vescia, The sequential hypothesis of the function of sleep. Behav. Brain Res. 69, 157–166 (1995).

9. S. Ribeiro, M. A. L. Nicolelis, Reverberation, storage, and postsynaptic propagation of memories during sleep. Learn. Mem. 11, 686–96 (2004).

10. P. A. Lewis, G. Knoblich, G. Poe, How Memory Replay in Sleep Boosts Creative Problem-Solving. Trends Cogn. Sci. 22, 491–503 (2018).

11. M. P. Walker, C. Liston, J. A. Hobson, R. Stickgold, Cognitive flexibility across the sleep-wake cycle: REM-sleep enhancement of anagram problem solving. Cogn. Brain Res. 14, 317–324 (2002).

12. R. Stickgold, L. Scott, C. Rittenhouse, J. a Hobson, Sleep-induced changes in associative memory. J. Cogn. Neurosci. 11, 182–193 (1999).

13. C. L. Edwards, P. M. Ruby, J. E. Malinowski, P. D. Bennett, M. T. Blagrove, Dreaming and insight. Front. Psychol. 4, 1–14 (2013).

14. F. Beijamini, S. I. R. Pereira, F. A. Cini, F. M. Louzada, After Being Challenged by a Video Game Problem, Sleep Increases the Chance to Solve It. PLoS One 9, e84342 (2014).

15. F. Beijamini, A. Valentin, R. Jäger, J. Born, S. Diekelmann, Sleep Facilitates Problem Solving With No Additional Gain Through Targeted Memory Reactivation. Front. Behav. Neurosci. 15, 30 (2021).

16. F. Ferrarelli, R. Kaskie, S. Laxminarayan, S. Ramakrishnan, J. Reifman, A. Germain, An increase in sleep slow waves predicts better working memory performance in healthy individuals. Neuroimage 191, 1–9 (2019).

17. K. A. Wilckens, F. Ferrarelli, M. P. Walker, D. J. Buysse, Slow-Wave Activity Enhancement to Improve Cognition. Trends Neurosci. 41, 470–482 (2018).

18. S. A. Cairney, A. á. V. Guttesen, N. El Marj, B. P. Staresina, Memory Consolidation Is Linked to Spindle-Mediated Information Processing during Sleep. Curr. Biol. 28, 948–954.e4 (2018).

19. L. M. J. Fernandez, A. L. Correspondence, A. Lüthi, X. Laura, M. J. Fernandez, Sleep spindles: Mechanisms and functions. Physiol. Rev. 100, 805–868 (2020).

20. R. Cox, W. F. Hofman, L. M. Talamini, Involvement of spindles in memory consolidation is slow wave sleep-specific. Learn. Mem. 19, 264–267 (2012).

21. B. E. Muehlroth, M. C. Sander, Y. Fandakova, T. H. Grandy, B. Rasch, Y. L. Shing, M. Werkle-Bergner, Precise Slow Oscillation–Spindle Coupling Promotes Memory Consolidation in Younger and Older Adults. Sci. Reports 2019 91 9, 1–15 (2019).

22. M. A. Hahn, D. Heib, M. Schabus, K. Hoedlmoser, R. F. Helfrich, Slow oscillation-spindle coupling predicts enhanced memory formation from childhood to adolescence. Elife 9, 1–21 (2020).

23. M. A. Wilson, B. L. McNaughton, Reactivation of Hippocampal Ensemble Memories During Sleep. Science (80-.). 265, 676–679 (1994).

24. D. B. Rubin, T. Hosman, J. N. Kelemen, A. Kapitonava, F. R. Willett, B. F. Coughlin, E. Halgren, E. Y. Kimchi, Z. M. Williams, J. D. Simeral, L. R. Hochberg, S. S. Cash, Learned Motor Patterns Are Replayed in Human Motor Cortex during Sleep. J. Neurosci. 42, 5007–5020 (2022).

25. M. Atienza, J. L. Cantero, Complex sound processing during human REM sleep by recovering information from long-term memory as revealed by the mismatch negativity (MMN). Brain Res. 901, 151–160 (2001).

26. E. J. Wamsley, R. Stickgold, Dreaming of a learning task is associated with enhanced memory consolidation: Replication in an overnight sleep study. J. Sleep Res. 28, e12749 (2019).

27. R. J. Katz, M. Steiner, Dream and motivation: a psychobiological approach. J. Soc. Biol. Struct. 2, 141–154 (1979).

28. C. Cavallero, P. Cicogna, V. Natale, M. Occhionero, A. Zito, Slow Wave Sleep Dreaming Sleep 15, 562–566 (1992).

29. A. Revonsuo, The reinterpretation of dreams: An evolutionary hypothesis of the function of dreaming. Behav. Brain Sci. 23, 877–901 (2000).

30. A. Revonsuo, K. Valli, Dreaming and Consciousness: Testing the Threat Simulation Theory of the Function of Dreaming. Psyche (Stuttg). 6 (2000).

31. K. Valli, A. Revonsuo, O. Pälkäs, K. H. Ismail, K. J. Ali, R. L. Punamäki, The threat simulation theory of the evolutionary function of dreaming: Evidence from dreams of traumatized children. Conscious. Cogn. 14, 188–218 (2005).

32. A. Zadra, S. Desjardins, É. Marcotte, Evolutionary function of dreams: A test of the threat simulation theory in recurrent dreams. Conscious. Cogn. 15, 450–463 (2006).

33. N. B. Mota, J. Weissheimer, M. Ribeiro, M. de Paiva, J. Avilla-Souza, G. Simabucuru, M. F. Chaves, L. Cecchi, J. Cirne, G. Cecchi, C. Rodrigues, M. Copelli, S. Ribeiro, Dreaming during the Covid-19 pandemic: Computational assessment of dream reports reveals mental suffering related to fear of contagion. PLoS One 15, e0242903 (2020).

34. K. Valli, A. Revonsuo, The threat simulation theory in light of recent empirical evidence: A review. Am. J. Psychol. 122, 17–38 (2009).

35. S. M. Fogel, L. B. Ray, V. Sergeeva, J. De Koninck, A. M. Owen, A novel approach to dream content analysis reveals links between learning-related dream incorporation and cognitive abilities. Front. Psychol. 9, 387144 (2018).

36. C. Picard-Deland, T. Aumont, A. Samson-Richer, T. Paquette, T. Nielsen, Whole-body procedural learning benefits from targeted memory reactivation in REM sleep and task-related dreaming. Neurobiol. Learn. Mem. 183, 107460 (2021).

37. S. F. Schoch, M. J. Cordi, M. Schredl, B. Rasch, The effect of dream report collection and dream incorporation on memory consolidation during sleep. J. Sleep Res. 28, e12754 (2019).

38. E. Werth, P. Achermann, A. A. Borbély, Fronto-occipital EEG power gradients in human sleep. J. Sleep Res. 6, 102–112 (1997).

39. G. Tinguely, L. A. Finelli, H. P. Landolt, A. A. Borbély, P. Achermann, Functional EEG topography in sleep and waking: State-dependent and state-independent features. Neuroimage 32, 283–292 (2006).

40. S. Ackermann, M. Cordi, R. La Marca, E. Seifritz, B. Rasch, Psychosocial Stress Before a Nap Increases Sleep Latency and Decreases Early Slow-Wave Activity. Front. Psychol. 10, 1–14 (2019).

41. J. Beck, E. Loretz, B. Rasch, Stress dynamically reduces sleep depth: temporal proximity to the stressor is crucial. Cereb. Cortex, doi: 10.1093/CERCOR/BHAC055 (2022).

42. A. K. Pesonen, T. Makkonen, M. Elovainio, R. Halonen, K. Räikkönen, L. Kuula, Presleep physiological stress is associated with a higher cortical arousal in sleep and more consolidated REM sleep. Stress 24, 667–675 (2021).

43. E. Hein, R. Halonen, T. Wolbers, T. Makkonen, M. Kyllönen, L. Kuula, I. Kurki, P. Stepnicka, A. K. Pesonen, Does sleep promote adaptation to acute stress: An experimental study. Neurobiol. Stress 29, 100613 (2024).

44. A. D. Krystal, J. D. Edinger, W. K. Wohlgemuth, G. R. Marsh, NREM Sleep EEG Frequency Spectral Correlates of Sleep Complaints in Primary Insomnia Subtypes. Sleep 25, 626–636 (2002).

45. N. Biabani, A. Birdseye, S. Higgins, A. Delogu, J. Rosenzweig, Z. Cvetkovic, A. Nesbitt, P. Drakatos, J. Steier, V. Kumari, D. O’Regan, I. Rosenzweig, The neurophysiologic landscape of the sleep onset: a systematic review. J. Thorac. Dis. 15, 4530–4543 (2023).

46. H. Merica, J. M. Gaillard, The EEG of the sleep onset period in insomnia: A discriminant analysis. Physiol. Behav. 52, 199–204 (1992).

47. L. Staner, F. Cornette, D. Maurice, G. Viardot, O. Le Bon, J. Haba, C. Staner, R. Luthringer, A. Muzet, J. P. Macher, Sleep microstructure around sleep onset differentiates major depressive insomnia from primary insomnia. J. Sleep Res. 12, 319–330 (2003).

48. K. Spiegelhalder, W. Regen, B. Feige, J. Holz, H. Piosczyk, C. Baglioni, D. Riemann, C. Nissen, Increased EEG sigma and beta power during NREM sleep in primary insomnia. Biol. Psychol. 91, 329–333 (2012).

49. M. L. Perlis, E. L. Kehr, M. T. Smith, P. J. Andrews, H. Orff, D. E. Giles, Temporal and stagewise distribution of high frequency EEG activity in patients with primary and secondary insomnia and in good sleeper controls. J. Sleep Res. 10, 93–104 (2001).

50. R. R. Freedman, EEG power spectra in sleep-onset insomnia. Electroencephalogr. Clin. Neurophysiol. 63, 408–413 (1986).

51. D. J. Buysse, A. Germain, M. L. Hall, D. E. Moul, E. A. Nofzinger, A. Begley, C. L. Ehlers, W. Thompson, D. J. Kupfer, EEG Spectral Analysis in Primary Insomnia: NREM Period Effects and Sex Differences. Sleep 31, 1673–1682 (2008).

52. M. Hall, J. F. Thayer, A. Germain, D. Moul, R. Vasko, M. Puhl, J. Miewald, D. J. Buysse, Psychological Stress Is Associated With Heightened Physiological Arousal During NREM Sleep in Primary Insomnia. Behav. Sleep Med. 5, 178–193 (2007).

53. M. Hall, D. J. Buysse, P. D. Nowell, E. A. Nofzinger, P. Houck, C. F. Reynolds, D. J. Kupfer, Symptoms of stress and depression as correlates of sleep in primary insomnia. Psychosom. Med. 62, 227–230 (2000).

54. D. Denis, S. Y. Kim, S. M. Kark, R. T. Daley, E. A. Kensinger, J. D. Payne, Slow oscillation-spindle coupling is negatively associated with emotional memory formation following stress. Eur. J. Neurosci. 55, 2632–2650 (2022).

55. G. S. Shields, M. A. Sazma, A. M. McCullough, A. P. Yonelinas, The effects of acute stress on episodic memory: A meta-analysis and integrative review. Psychol. Bull. 143, 636–675 (2017).

56. J. D. Payne, E. D. Jackson, S. Hoscheidt, L. Ryan, W. J. Jacobs, L. Nadel, Stress administered prior to encoding impairs neutral but enhances emotional long-term episodic memories. Learn. Mem. 14, 861–868 (2007).

57. T. J. Cunningham, S. L. Leal, M. A. Yassa, J. D. Payne, Post-encoding stress enhances mnemonic discrimination of negative stimuli. Learn. Mem. 25, 611–619 (2018).

58. T. T. Dang-Vu, A. Salimi, S. Boucetta, K. Wenzel, J. O’Byrne, M. Brandewinder, C. Berthomier, J. P. Gouin, Sleep spindles predict stress-related increases in sleep disturbances. Front. Hum. Neurosci. 9, 128426 (2015).

59. S. Diekelmann, S. Biggel, B. Rasch, J. Born, Offline consolidation of memory varies with time in slow wave sleep and can be accelerated by cuing memory reactivations. Neurobiol. Learn. Mem. 98, 103–111 (2012).

60. M. Strauss, L. Griffon, P. Van Beers, M. Elbaz, J. Bouziotis, F. Sauvet, M. Chennaoui, D. Léger, P. Peigneux, Order matters: sleep spindles contribute to memory consolidation only when followed by rapid-eye-movement sleep. Sleep 45 (2022).

61. A. C. Souza, D. C. F. Golbert, J. S. Cassoli, I. Sánchez-Gendriz, V. V. F. Lima, F. A. Cini, D. Martins-de-Souza, S. Ribeiro, Experience-dependent phosphoproteomic changes in hippocampus and neocortex correlate with the abundance of spindle oscillations during the transition between SWS and REM sleep. doi: 10.21203/RS.3.RS-1842393/V2 (2022).

62. W. Blanco, C. M. Pereira, V. R. Cota, A. C. Souza, C. Rennó-Costa, S. Santos, G. Dias, M. G. Guerreiro, A. B. L. Tort, A. ãD Neto, S. Ribeiro, Synaptic Homeostasis and Restructuring across the Sleep-Wake Cycle. PLOS Comput. Biol. 11, e1004241 (2015).

63. R. Stickgold, A. Malia, D. Maguire, D. Roddenberry, M. O’Connor, Replaying the game: Hypnagogic images in normals and amnesics. Science (80-.). 290, 350–353 (2000).

64. C. Kusse, A. Shaffii-Le Bourdiec, J. Schrouff, L. Matarazzo, P. Maquet, Experience-dependent induction of hypnagogic images during daytime naps: a combined behavioural and EEG study. J. Sleep Res. 21, 10–20 (2012).

65. E. J. Wamsley, K. Perry, I. Djonlagic, L. B. Reaven, R. Stickgold, Cognitive Replay of Visuomotor Learning at Sleep Onset: Temporal Dynamics and Relationship to Task Performance. Sleep 33, 59–68 (2010).

66. E. J. Wamsley, M. Tucker, J. D. Payne, J. A. Benavides, R. Stickgold, Dreaming of a Learning Task Is Associated with Enhanced Sleep-Dependent Memory Consolidation. Curr. Biol. 20, 850–855 (2010).

67. A. W. Stamm, N. D. Nguyen, B. J. Seicol, A. Fagan, A. Oh, M. Drumm, M. Lundt, R. Stickgold, E. J. Wamsley, Negative reinforcement impairs overnight memory consolidation. Learn. Mem. 21, 591–596 (2014).

68. L. Hudachek, E. J. Wamsley, A meta-analysis of the relation between dream content and memory consolidation. Sleep 46 (2023).

69. E. J. Wamsley, K. Hamilton, Y. Graveline, S. Manceor, E. Parr, Test Expectation Enhances Memory Consolidation across Both Sleep and Wake. PLoS One 11, e0165141 (2016).

70. J. Pan, J. Wu, J. Liu, J. Wu, F. Wang, A Systematic Review of Sleep in Patients with Disorders of Consciousness: From Diagnosis to Prognosis. Brain Sci. 2021, 11, 1072. mdpi.com, doi: 10.3390/brainsci (2021).

71. A. N. Bertolazi, S. C. Fagondes, L. S. Hoff, E. G. Dartora, I. C. da Silva Miozzo, M. E. F. de Barba, S. S. Menna Barreto, Validation of the Brazilian Portuguese version of the Pittsburgh Sleep Quality Index. Sleep Med. 12, 70–75 (2011).

72. S. M. G. P. Togeiro, A. K. Smith, Métodos diagnósticos nos distúrbios do sono. Rev. Bras. Psiquiatr. 27, 8–15 (2005).

73. A. N. Bertolazi, S. C. Fagondes, L. S. Hoff, V. D. Pedro, S. S. Menna Barreto, M. W. Johns, Portuguese-language version of the Epworth sleepiness scale: validation for use in Brazil. J. Bras. Pneumol. 35, 877–883 (2009).

74. V. Jurcak, D. Tsuzuki, I. Dan, 10/20, 10/10, and 10/5 systems revisited: their validity as relative head-surface-based positioning systems. Neuroimage 34, 1600–1611 (2007).

75. A. Delorme, S. Makeig, EEGLAB: An open source toolbox for analysis of single-trial EEG dynamics including independent component analysis. J. Neurosci. Methods 134, 9–21 (2004).

76. L. J. Gabard-durnam, A. S. M. Leal, C. L. Wilkinson, The Harvard Automated Processing Pipeline for Electroencephalography (HAPPE): Standardized Processing Software for Developmental and High-Artifact Data. 12, 1–24 (2018).

77. L. Pion-Tonachini, K. Kreutz-Delgado, S. Makeig, ICLabel: An automated electroencephalographic independent component classifier, dataset, and website. Neuroimage 198, 181–197 (2019).

78. R. Debnath, G. A. Buzzell, S. Morales, M. E. Bowers, S. C. Leach, N. A. Fox, The Maryland analysis of developmental EEG (MADE) pipeline. Psychophysiology 57, e13580 (2020).

79. E. M. Whitham, K. J. Pope, S. P. Fitzgibbon, T. Lewis, C. R. Clark, S. Loveless, M. Broberg, A. Wallace, D. DeLosAngeles, P. Lillie, A. Hardy, R. Fronsko, A. Pulbrook, J. O. Willoughby, Scalp electrical recording during paralysis: Quantitative evidence that EEG frequencies above 20 Hz are contaminated by EMG. Clin. Neurophysiol. 118, 1877–1888 (2007).

80. D.-W. Kim, C.-H. Im, “EEG Spectral Analysis” in Computational EEG Analysis: Methods and Applications, C.-H. Im, Ed. (Springer Singapore, Singapore, 2018; http://www.springer.com/series/3740) Biological and Medical Physics, Biomedical Engineering, pp. 35–55.

81. K. Lacourse, J. Delfrate, J. Beaudry, P. Peppard, S. C. Warby, A sleep spindle detection algorithm that emulates human expert spindle scoring. J. Neurosci. Methods 316, 3–11 (2019).

82. D. T. Plante, M. R. Goldstein, J. D. Cook, R. Smith, B. A. Riedner, M. E. Rumble, L. Jelenchick, A. Roth, G. Tononi, R. M. Benca, M. J. Peterson, Effects of partial sleep deprivation on slow waves during non-rapid eye movement sleep: A high density EEG investigation. Clin. Neurophysiol. 127, 1436–1444 (2016).

83. J. Pan, W. J. Tompkins, A Real-Time QRS Detection Algorithm. IEEE Trans. Biomed. Eng. BME-32, 230–236 (1985).

84. J. Ramshur, Design, Evaluation, and Application of Heart Rate Variability Analysis Software (HRVAS). Electron. Theses Diss. (2010).

85. F. Ritsert, M. Elgendi, V. Galli, C. Menon, Heart and Breathing Rate Variations as Biomarkers for Anxiety Detection. Bioengineering 9, 711 (2022).

86. M. A. Jaoude, H. Sun, K. R. Pellerin, M. Pavlova, R. A. Sarkis, S. S. Cash, M. Brandon Westover, A. D. Lam, Expert-level automated sleep staging of long-term scalp electroencephalography recordings using deep learning. Sleep 43 (2020).

87. T. K. Koo, M. Y. Li, A Guideline of Selecting and Reporting Intraclass Correlation Coefficients for Reliability Research. J. Chiropr. Med. 15, 155 (2016).

88. D. M. Groppe, T. P. Urbach, M. Kutas, Mass univariate analysis of event-related brain potentials/fields II: Simulation studies. Psychophysiology 48, 1726–1737 (2011).

