## Supplemental Tables and Texts for "Dream content and slow waves benefit preys against predators in a video game confrontation"

Daniel Brandão et. al.

**The PDF file includes:**

Tables S1 to S2

Supplementary Text

***Table S1.***

*A priori comparison between participants in the prey and predator groups*

| Variable | Predator | Prey | p-value | p-sig |
|---|---|---|---|---|
| **Sex (M/F)** | 11/2 | 12/1 | 0,5393 | n.s. |
| **Gaming Experience (0/1)** | 3/10 | 2/11 | 0,6188 | n.s. |
| **FPS Experience (0/1)** | 2/11 | 2/11 | 1,0000 | n.s. |
| **Interest in FPS (0/1/2)** | 2/5/6 | 2/4/7 | 0,9103 | n.s. |
| **Age** | 23,9231 | 25,3077 | 0,2208 | n.s. |
| **Total sleep time last night (h)** | 6,0769 | 6,1538 | 0,5334 | n.s. |
| **Epworth (Sleepness)** | 7,6923 | 7,6923 | 1,0000 | n.s. |
| **Sleep quality** | 1,0769 | 1,0000 | 0,7729 | n.s. |
| **Sleep latency** | 2,3846 | 1,4615 | 0,0515 | n.s. |
| **Sleep duration** | 0,3846 | 0,3077 | 0,7909 | n.s. |
| **Sleep efficiency** | 0,5385 | 0,0769 | 0,1355 | n.s. |
| **Sleep disorders** | 0,9231 | 1,0769 | 0,1831 | n.s. |
| **Sleep medication** | 0,0000 | 0,1538 | 0,3560 | n.s. |
| **Daytime dysfunction** | 1,1538 | 1,2308 | 0,9771 | n.s. |
| **Pittsburgh Total** | 6,4615 | 5,3077 | 0,3789 | n.s. |

***Table S2.***

*A priori comparison between the two samples of study participants*

| Variable | Sample 1 | Sample 2 | p-value | p-sig |
|---|---|---|---|---|
| **Sex (M/F)** | 15/3 | 8/0 | 0,2196 | n.s. |
| **Gaming Experience (0/1)** | 3/15 | 2/6 | 0,6188 | n.s. |
| **FPS Experience (0/1)** | 3/15 | 1/7 | 0,7858 | n.s. |
| **Interest in FPS (0/1/2)** | 4/6/8 | 0/3/5 | 0,3385 | n.s. |
| **Age** | 24,7222 | 24,3750 | 0,7349 | n.s. |
| **Total sleep time last night (h)** | 6,6944 | 4,8125 | 0,1154 | n.s. |
| **Epworth (Sleepness)** | 7,9444 | 7,1250 | 0,5200 | n.s. |
| **Sleep quality** | 1,0000 | 1,1250 | 0,6845 | n.s. |
| **Sleep latency** | 1,6111 | 2,6250 | 0,1204 | n.s. |
| **Sleep duration** | 0,3889 | 0,2500 | 0,8575 | n.s. |
| **Sleep efficiency** | 0,3333 | 0,2500 | 0,7772 | n.s. |
| **Sleep disorders** | 1,0000 | 1,0000 | 1,0000 | n.s. |
| **Sleep medication** | 0,1111 | 0,0000 | 0,5597 | n.s. |
| **Daytime dysfunction** | 1,1667 | 1,2500 | 0,9010 | n.s. |
| **Pittsburgh Total** | 5,6111 | 6,5000 | 0,3848 | n.s. |

**Supplementary Text**

Representative example of a dream report highly related with the game (rating = 4.5)

Well, I dreamed about several things. First, what I remember what I dreamed about, was actually about the experiment I was doing here. I dreamed that the researcher was talking to me about the experiment and that there were other types of chairs here in the experiment. I remember there were some chairs here. Then I dreamed that my girlfriend was here too and she stayed here in the experiment and watching how me and the other player were doing. Afterwards I dreamed about the game. I dreamed about playing the game again.

Representative example of a dream report highly unrelated with the game (rating = 1.5)

I was dreaming that I was on a beach. There were a lot of people and I was with my dog until it rained and then I started dreaming about my grandparents
